# TGS1: a novel regulator of β-cell mass and function

**DOI:** 10.1101/2021.09.20.461105

**Authors:** Manuel Blandino-Rosano, Pau Romaguera Llacer, Ashley Lin, Janardan K Reddy, Ernesto Bernal-Mizrachi

## Abstract

Type 2 diabetes (T2D) is a metabolic disorder associated with abnormal glucose homeostasis and is characterized by intrinsic defects in β-cell function and mass. Trimethylguanosine synthase 1 (TGS1) is an evolutionarily conserved enzyme that methylates small nuclear and nucleolar RNAs (snRNAs and snoRNAs) and is involved in pre-mRNA splicing, transcription, and ribosome production. However, the role of TGS1 in β-cells and glucose homeostasis had not been explored. Here we show that TGS1 is upregulated by insulin and upregulated in islets from mice exposed to a high-fat diet and in human β-cells from T2D donors. Using mice with conditional (*βTGS1KO* and *βTGS1^Het^)* and inducible (*MIP-Cre^ERT^-TGS1KO*) TGS1 deletion, we determine that TGS1 regulates β-cell mass and function. Unbiased approaches allowed us to identify a link between TGS1 and ER stress and cell cycle arrest and how TGS1 regulates β-cell apoptosis. Deletion of TGS1 results in an increase in the unfolded protein response by increasing XBP-1, ATF-4, and the phosphorylation of eIF2α, and several changes in cell cycle inhibitors and activators such as p27 and Cyclin D2. This study establishes TGS1 as a key player regulating β-cell mass and function as well as playing a role in the adaptive β-cell function to a high-fat diet. These observations can be used as a stepping-stone for the design of novel strategies using TGS1 as a therapeutic target for the treatment of diabetes.

## INTRODUCTION

Type 2 diabetes (T2D) is a disease of epidemic proportions and a major public health problem with a total estimated cost of $245 billion (1–5). T2D is characterized by insulin resistance at peripheral tissues such as skeletal muscle and liver, and intrinsic defects in β-cell function and mass. β-cells primarily adapt to insulin resistance by increasing function and proliferation, although cell size and apoptosis also play a role (6). However, a decompensated phase is observed when β-cells are not able to respond to the demands of insulin required to control glucose homeostasis, resulting in impaired function and deterioration of β-cells mass. Despite several mechanisms have been proposed, it is believed that chronic exposure of β-cells to hyperglycemia alters the protein folding capacity of the endoplasmic reticulum (ER), resulting in the accumulation and aggregation of unfolded proteins causing ER stress (7).

TGS1 is an evolutionarily conserved enzyme that mediates methylation of small nuclear and nucleolar RNAs (snRNAs and snoRNAs), selenoprotein and telomerase mRNAs and is involved in pre-mRNA splicing, transcription, and ribosome biogenesis (8,9). Initially, snRNAs and snoRNAs are capped with 7-monomethylguanosine (m^7^G) and methylated by TGS1 to form 2,2,7-trimethylguanosine (TMG). The TMG cap modification is highly conserved throughout eukaryotes and is a critical requirement for RNA transport and regulation of splicing (10). More importantly, recent studies have demonstrated that impaired snRNPs biogenesis affects glucose metabolism and pancreatic development in mouse and human (11). While very little is known about TGS1 in vertebrates and virtually nothing about TGS1 in β-cells, previous studies indicate that TGS1 could be involved in controlling glucose homeostasis (11–15). Mice with a conditional deletion of TGS1 in liver exhibited alterations in liver-specific metabolic pathways followed by impaired hepatic gluconeogenesis (15). In addition, higher TGS1 levels in the soleus muscle of rats exposed to high sucrose diet-induced insulin resistance was previously observed (14). More importantly, TGS1 was elevated in cultured myoblasts stimulated with insulin and TGS1 overexpression in rat skeletal muscle and cultured myoblasts inhibited insulin-stimulated glucose uptake, further supporting the role of TGS1 in regulation of insulin sensitivity (14). These findings pointed towards an important role of TGS1 in regulating hepatic gluconeogenesis, insulin resistance and glucose homeostasis. On the other hand, the role of TGS1 on pancreatic β-cell function is completely unknown.

Given the important role of snRNAs and snoRNAs in β-cells and pancreas development (11,12) and the association between TGS1 upregulation in hyperglycemia and regulation of hepatic glucose output (14,15), we decided to study TGS1 in β-cells. Herein we observed that TGS1 levels are increased in high-fat fed (HFD) mice and humans with T2D. Using mice with conditional TGS1 deletion in β-cells (*βTGS1KO*), the current studies show that TGS1 deficiency results in decreased β-cell function and mass due to increased ER stress, apoptosis, and cell cycle arrest. RNA-Seq analysis in control and *βTGS1KO* islets at 1 month of age demonstrated differential expression in several ER stress and cell cycle arrest genes. Despite heterozygous TGS1 apparently did not show any changes in glucose handling on regular chow, impairment of glucose homeostasis following HFD demonstrates that the maintenance of narrow levels of TGS1 is indispensable for β-cell adaptation to insulin resistance.

## RESULTS

### TGS1 levels are elevated in models of diabetes and induced by glucose and insulin in a paracrine/autocrine manner

To study the regulation of TGS1 by glucose in β-cells, isolated wild-type islets were treated with 3 or 16mM glucose for 24h. High glucose exposure increased protein levels of TGS1 by immunostaining (Fig. 1a). To test the paracrine role of insulin on TSG1 upregulation by glucose, we used Somatostatin-28, a potent inhibitor of insulin secretion. Somatostatin-28 treatment prevented the TSG1 upregulation induced by high glucose, indicating that high glucose induced TGS1 expression by insulin in a paracrine/autocrine fashion (Fig. 1a). To test the direct effect of insulin, we treated wild-type islets with low glucose and insulin (100 nM) for 24h. TGS1 staining and immunoblotting show that insulin treatment increased TGS1 levels in islets from WT mice (Figs. 1b-c). These data suggest that insulin, but not glucose, regulates TGS1 expression. Consistent with these data, TGS1 levels in β-cells are elevated in conditions of HFD induced insulin resistance and hyperglycemia (mice fed HFD for 8 weeks) (Fig. 1d). Assessment of TGS1 expression in purified β-cells (reanalysis of a previously published RNA-Seq data set (E-MTAB-5061/5060) (16)) and stained pancreatic sections from human donors showed that TGS1 mRNA and protein were increased in T2D donors compared to controls (Figs. 1e-f). Moreover, the extent to which this increase in TGS1 plays either a beneficial compensatory role or induces β-cell failure was unexplored.

**Figure 1.**
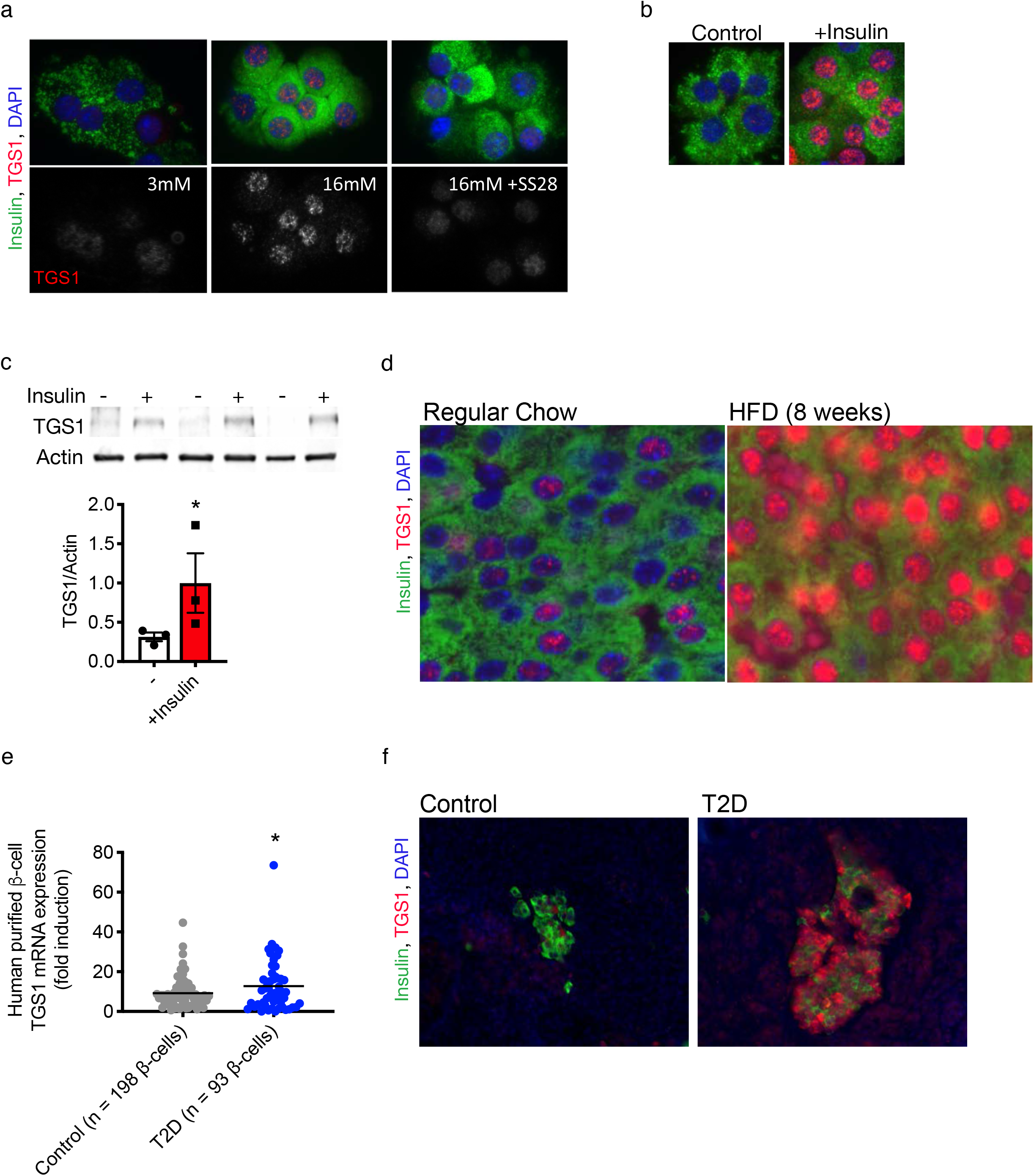
Insulin regulates TGS1 expression. (a) Immunostaining of insulin (green), TGS1 (red, upper, and white, lower) and DAPI (blue) in dispersed islets from WT mice exposed to 3, 16mM and 16mM glucose +SS28 for 24h. (b) Immunostaining of insulin (green), TGS1 (red) and DAPI (blue) of dispersed β-cells from WT islets treated with or without insulin (100nM) for 24h. (c) Immunoblotting and quantification of TGS1 and actin in WT islets treated with or without insulin (100nM) at 3mM glucose for 24h (n=3). (d) Immunostaining for insulin (green), TGS1 (red) and DAPI (blue) in pancreas sections from mice on regular chow or HFD. (e) RNA-Seq dataset from healthy (n=198) and T2D (n=93) purified human β-cells obtained from public repositories (E-MTAB-5061/5060) (16). (f) Immunostaining for insulin (green), TGS1 (red) and DAPI (blue) in pancreas sections from control and T2D donors. Stainings show a representative image from three independent experiments. Data expressed as means±s.e.m., \**p*<0.05 compared to samples control within the group assessed by one-way ANOVA.

### Generation of mice with TGS1 inactivation in β-cells

To inactivate TGS1 function, we generated mice with homozygous deletion of TGS1 in β-cells by crossing *TGS1^f/f^* (17) with *RIP-Cre* mouse (18) (*βTGS1KO*). Figure 2a shows that TGS1 is absent in β-cells from *βTGS1KO* islets, and this was confirmed by immunoblotting (Fig. 2b). The 2,2,7-trimethylguanosine (TMG) cap is synthesized by TGS1 and is abundantly present on snRNAs essential for pre-mRNA splicing (8,9,19–24). Previous studies have shown that the TMG cap provides a signal for the efficient nuclear import of the newly assembled snRNPs (8,19,25,26). To validate the inactivation of TGS1 in β-cells, we assessed U1 snRNP nuclear levels. Immunostaining of pancreas sections from control and *βTGS1KO* mice showed a 30% reduction of nuclear U1 snRNP levels with no differences in total U1 snRNP levels in β-cells from *βTGS1KO* mice (Figs. 2c-d). Altered snRNPs translocation to the nucleus has been associated with disruption in Cajal Bodies (CBs) formation (27,28). To examine CB formation, we assessed coilin levels, a well-known marker of CBs. Coilin levels were decreased, and this was associated with reduced numbers of CBs (Figs. 2e and f). Taken together, these experiments validate the inactivation of TGS1 function in *βTGS1KO* mice.

**Figure 2.**
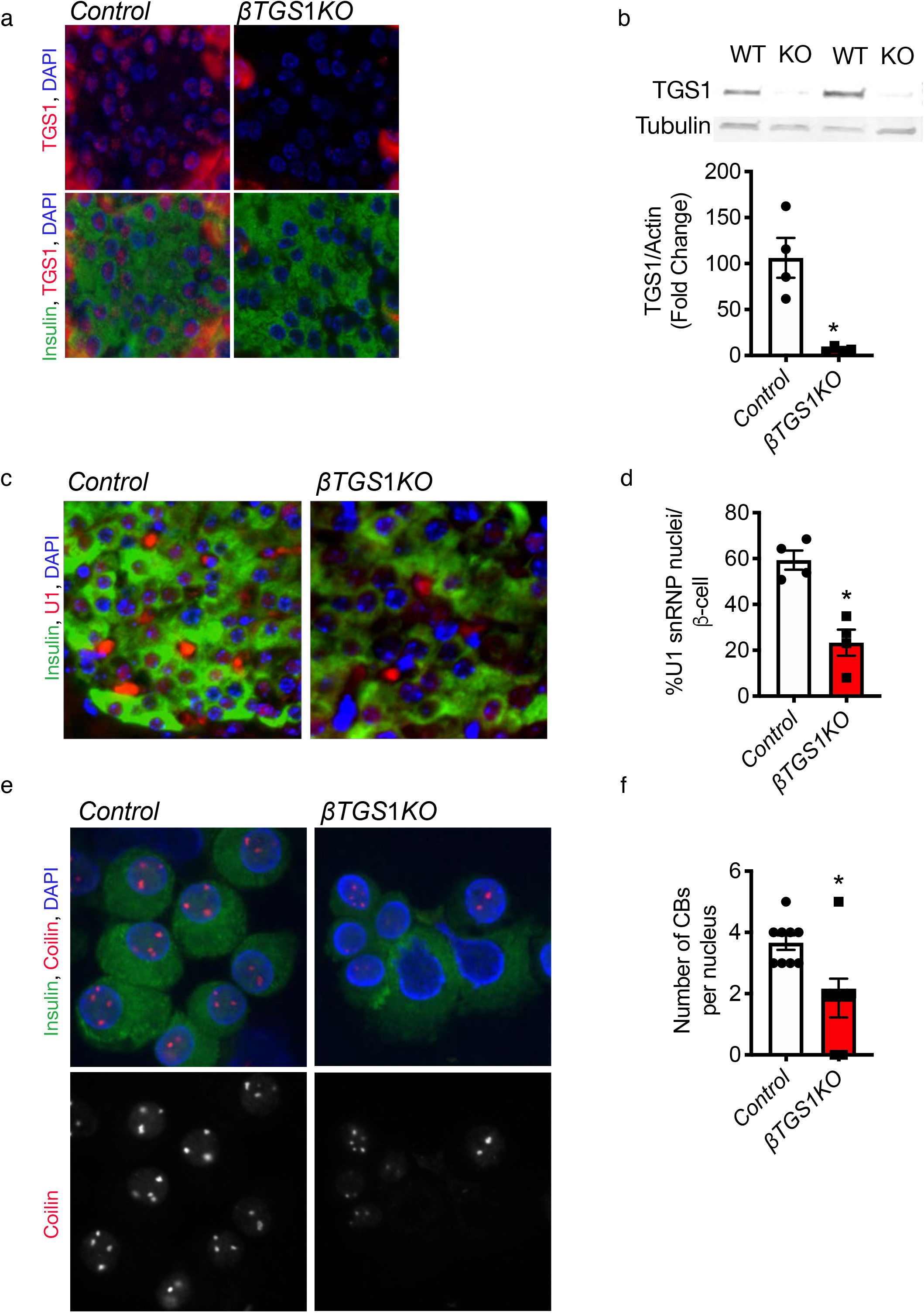
TGS1 deletion decreased nuclear levels of U1 and coilin and reduced Cajal Bodies number. (a) Immunostaining for TGS1 (red), insulin (green) and DAPI (blue) in pancreas sections from *control* and *βTGS1KO.* (b) Immunoblotting and quantification of TGS1 and tubulin in isolated islets from control and *βTGS1KO*. Staining (c) and quantification (d) of nuclear U1 (red), insulin (green) and DAPI (blue) in pancreas sections from *control* and *βTGS1KO* mice. (e) Staining for coilin (red, upper and white, lower) in dispersed islets from control and *βTGS1KO* mice. (f) Number of CBs per nucleus in *βTGS1KO* and controls (n=4). Staining shows a representative image from four independent experiments. Data expressed as means±s.e.m. \**p*<0.05. Nonparametric *U*-test (Mann–Whitney).

### Constitutive deletion of TGS1 in β-cells results in hyperglycemia

*βTGS1KO* mice exhibit a normal response to glucose and normal fed glucose and insulin levels up to 1 month of age (Figs. 3a-c). Hypoinsulinemia and hyperglycemia in the fed state were observed in *βTGS1KO* mice after 1 month of age (Figs. 3a, and d-g). At 3 months, male *βTGS1KO* mice exhibit an increase in fed glucose (Fig. 3a,) decrease in insulin (Fig. 3d) and impaired glucose tolerance after glucose challenge (Fig. 3e). This phenotype was also observed in females, suggesting that sexual dimorphism is not playing a role in the TGS1 deficient mice (Fig. 3f-g). Moreover, *in vitro* glucose-stimulated insulin secretion was blunted in islets from *βTGS1KO* mice at 1 month of age and this is not explained by alterations in insulin content (Figs. 3h and i). To validate the importance of TGS1 in insulin secretion, we deleted TGS1 in two-month-old mice using a mouse model with tamoxifen (TMX) inducible deletion of TGS1 in β-cells (*MIP-Cre^ERT^-TGS1KO*). These studies show that *MIP-Cre^ERT^-TGS1KO* mice display elevation in fed glucose at 3 weeks post-TMX injection (Figs. 3j-k). In addition, glucose tolerance was impaired in *MIP-Cre^ERT^-TGS1KO* mice, and this was accompanied by comparable β-cell mass (Figs. 3l-m).

**Figure 3.**
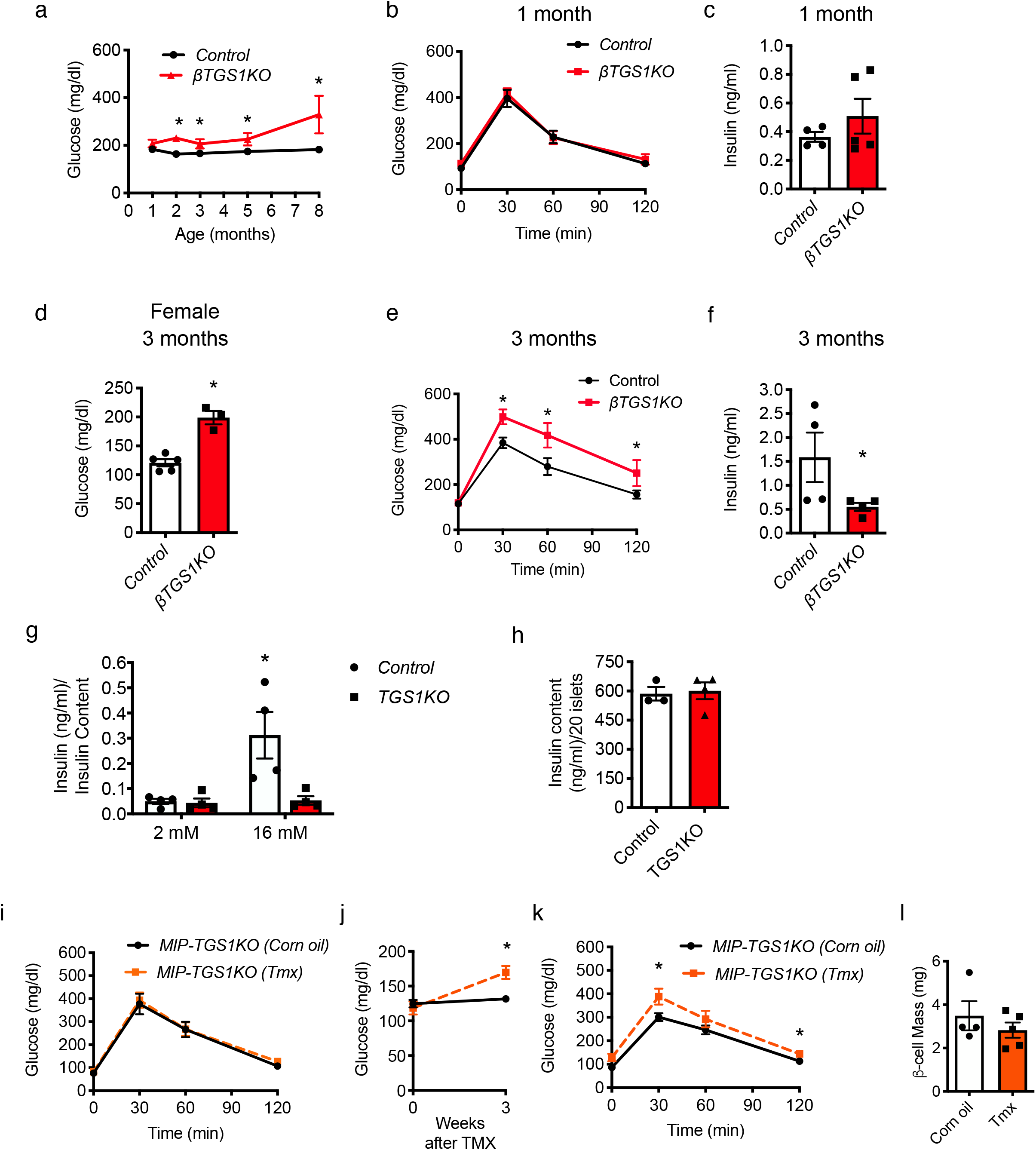
Constitutive deletion of TGS1 in β-cells results in hyperglycemia. (a) Random fed glucose levels during the first 8 months of age. (b) Random fed serum insulin levels at 1 month of age. (c) Intraperitoneal glucose tolerance test (IPGTT) at 1 month of age. (d) Random fed serum insulin levels at 3 months of age. (e) IPGTT at 3 months of age. (f) Random fed glucose levels in females at 3 months of age. (g) IPGTT in females at 3-4 months of age. (h) Glucose stimulated insulin secretion (GSIS) *in vitro* using isolated islets from 2-month-old. (i) Insulin content. (n=4). (j) IPGTT before TMX injection at 1 months of age. (k) Random fed glucose levels before and 3 weeks post-TMX injection. (l) IPGTT 3 weeks post-TMX injection in *MIP-TGS1KO*. (m) Assessment of β-cell mass 4 weeks after TMX injection. Data expressed as means±s.e.m., \**p*<0.05. Nonparametric *U*-test (Mann–Whitney) and \**p*<0.05 compared to 2m glucose within the group assessed by two-way ANOVA.

### βTGS1KO exhibits a reduction in β-cell mass

To determine if the defect in insulin secretion results from alterations in stimulus-secretion coupling and/or a reduction in β-cell mass, we performed assessment of β-cell morphometry. β-cell mass in *βTGS1KO* mice was reduced at 3 months of age (Figs. 4a-b). Decreased in β-cell mass in *βTGS1KO* mice was associated with increased in β-cell apoptosis assessed by TUNEL and active Caspase 3 measured by FACS (Fig. 4 c-d). This was accompanied by increased in β-cell proliferation assessed by Ki67 staining (Fig. 4e). No changes in β-cell size were observed in *βTGS1KO* mice (Figs. 4f).

**Figure 4.**
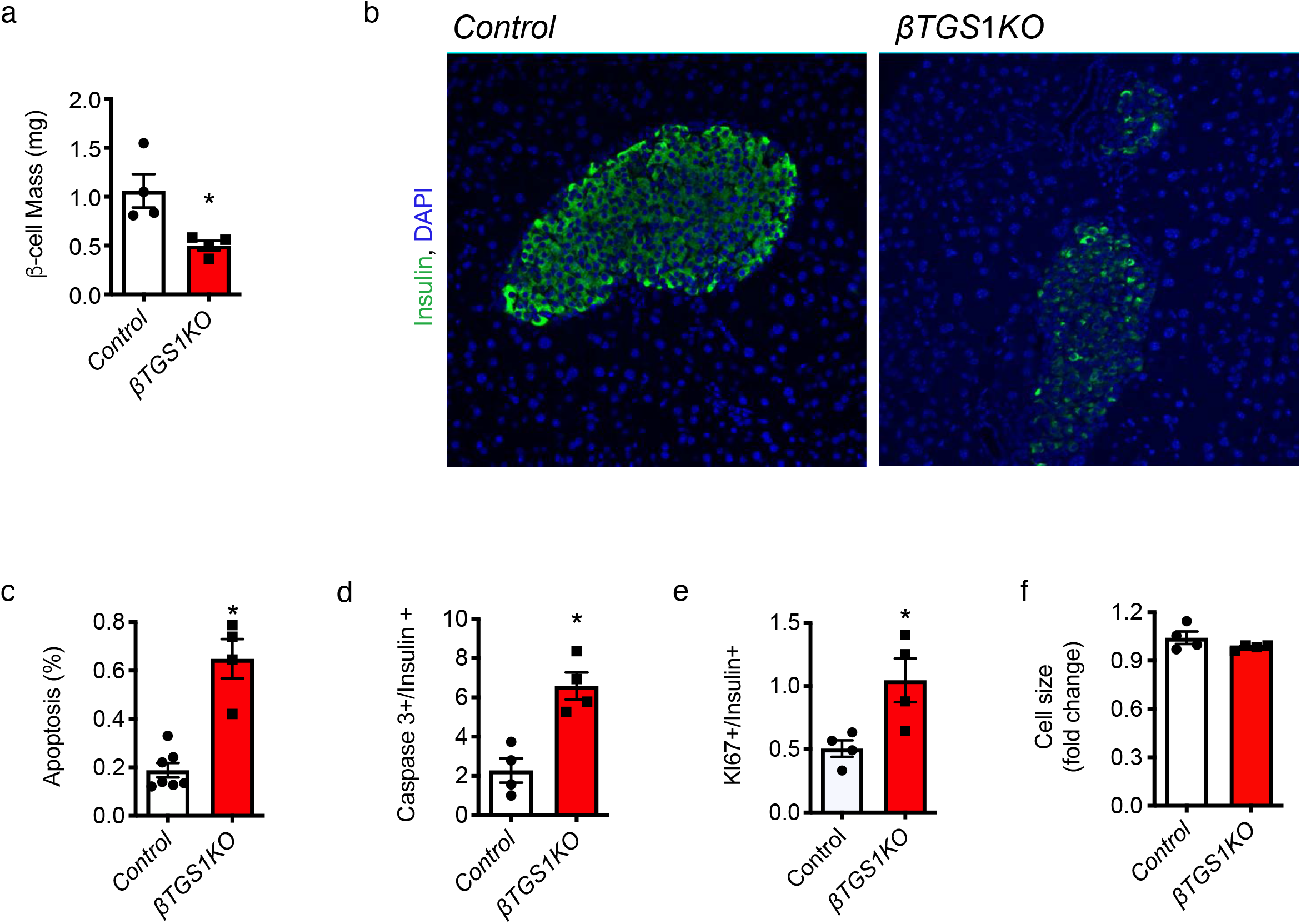
*βTGS1KO* mouse exhibits a reduction in β-cell mass due to increase in apoptosis at 3 months of age. (a) Assessments of β-cell mass and (b) representative pictures of islets from *βTGS1KO* and control mice, (c) apoptosis in β-cells by TUNEL and (d) Caspase 3 positive in insulin positive cells by FACS, (e) proliferation by Ki67 staining and (f) cell size. Data expressed as means±s.e.m., \**p*<0.05. Nonparametric *U*-test (Mann–Whitney).

### RNA-Seq studies in βTGS1KO reveal alterations in markers of ER stress

To test how TGS1 regulates β-cell apoptosis, we performed RNA-Seq in islets from *βTGS1KO* before the development of hyperglycemia and reduction of β-cell mass (1 month-old). We used EnrichR (29) to perform gene ontology (GO)-term analysis on the full list of genes differentially expressed. GO-term analysis revealed increased expression patterns (>1.5-fold change) in genes involved in protein transport, insulin, protein hormone and protein secretion and ER stress (Fig. 5a). Decreased expression (<0.5-fold change) of genes associated with cellular processes including cell migration, proliferation, migration and extracellular matrix organization (Fig. 5b). Unbiased pathway enrichment analysis indicated that genes related to unfolded protein response (UPR) were increased in *βTGS1KO* (z-score=5.61). The mRNA of BIP (hspa5), a protein that interacts with the three UPR activator proteins, PERK, ATF6, and IRE1α acting as a repressor of the UPR and playing a role in the apoptosis process was significantly increased in *βTGS1KO* islets (Fig. 5c). It was not unexpected to see that genes related with the three separate branches of the ER response were also increased: 1. PERK (eif2ak3), atf4, ddit3 (CHOP/GADD153) and Sestrins), 2. ATF6, and 3. IRE1α (ern1, gadd34, trib3, and tspyl2). In addition, other top differentially expressed genes related to ER response, like wfs1, dnajb9 or hyou1 are also observed in Fig. 5c. Given the importance of ER stress in β-cells, we validated the abnormalities in UPR pathway gene expression by immunoblotting. Figure 5d shows a 2-fold increase in XBP-1 confirming the activation of UPR pathways via the induction of PERK. eIF2α phosphorylation and ATF4 were elevated in islets from *βTGS1KO* (Fig. 5d). Electron microscopy shows that ER of β-cells from *βTGS1KO* mice was enlarged (Fig. 5e).

**Figure 5.**
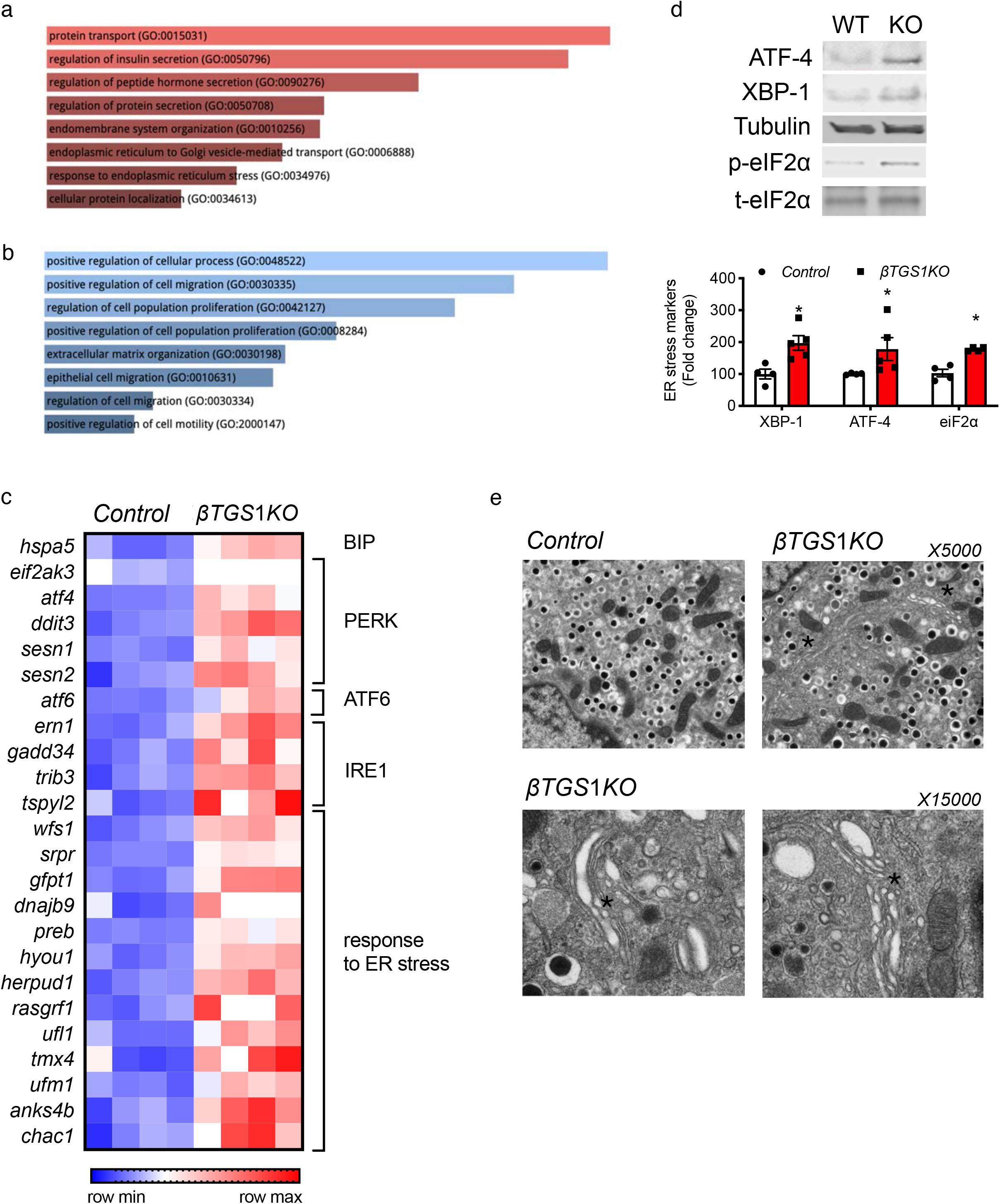
*β*-*cells with deletion of TGS1* exhibit signs of ER stress. GO-driven pathway analysis of differentially expressed genes with (a) > 1.5-fold change and (b) <0.5-fold change between *βTGS1KO* and control islets at 1 month of age. (c) Gene-expression heatmap of the differentially expressed genes related to unfolded protein response (UPR) and the response to ER stress in *βTGS1KO* islets compared to control. Genes are represented in rows and mice in columns. (d) Immunoblotting and quantification of ATF-4, XBP-1 respect to tubulin and phospho respect to total eiF2α in islets from *control* and *βTGS1KO* mice at 1 month of age. (e) Electron microscopy images showing enlarged ER in β-cells from *βTGS1KO* mice at 1 month of age. Magnification: ×5000 and x15000. Images are representative of islets from three animals. Asterisks indicate areas showing ER enlarged. Data expressed as means±s.e.m. \**p*<0.05. Nonparametric *U*-test (Mann–Whitney).

### βTGS1KO exhibits abnormalities in cell cycle progression

The morphometric analysis suggested that the increase in proliferation was not accompanied by increases in β-cell mass indicating that cell cycle progression abnormalities could be involved. To assess this, we compared the results of Ki67 (marker of all phases of the cell cycle, Fig. 4e) with bromodeoxyuridine (BrdU) (S phase). The fraction of BrdU in β-cells was similar between *βTGS1KO* and control mice (Fig. 6b). To assess the mechanisms of cell cycle abnormalities, we analysed the RNA-Seq data. mRNAs of G1-S cell cycle inhibitors p18, p19, and p27 were increased in *βTGS1KO* islets (Figs. 5b and 6a). Analysis of mRNAs of cell cycle components that drive G1-S transition showed increase in cyclins D1 and D2, necessary regulators for cell cycle entry but significantly decreased levels of cyclin D partners cdk6, cdk4 in *βTGS1KO* mice. Other G1-S transition genes, cyclin E and cdk2, were also reduced in *βTGS1KO* mice (Fig. 6a). Cyclin A, cyclin B and cdk1 mRNAs, genes implicated in S, G2 and M phase, were also decreased in *βTGS1KO* mice (Fig. 6a). The increases in Cyclin D2 and p27 mRNAs, two key regulators of G1-S transition in β-cells, were validated by immunoblotting (Figs. 6c). Taken together, these experiments suggest that deletion of TGS1 in β-cells entered the cell cycle but exhibited abnormal transition and completion of the cell cycle.

**Figure 6.**
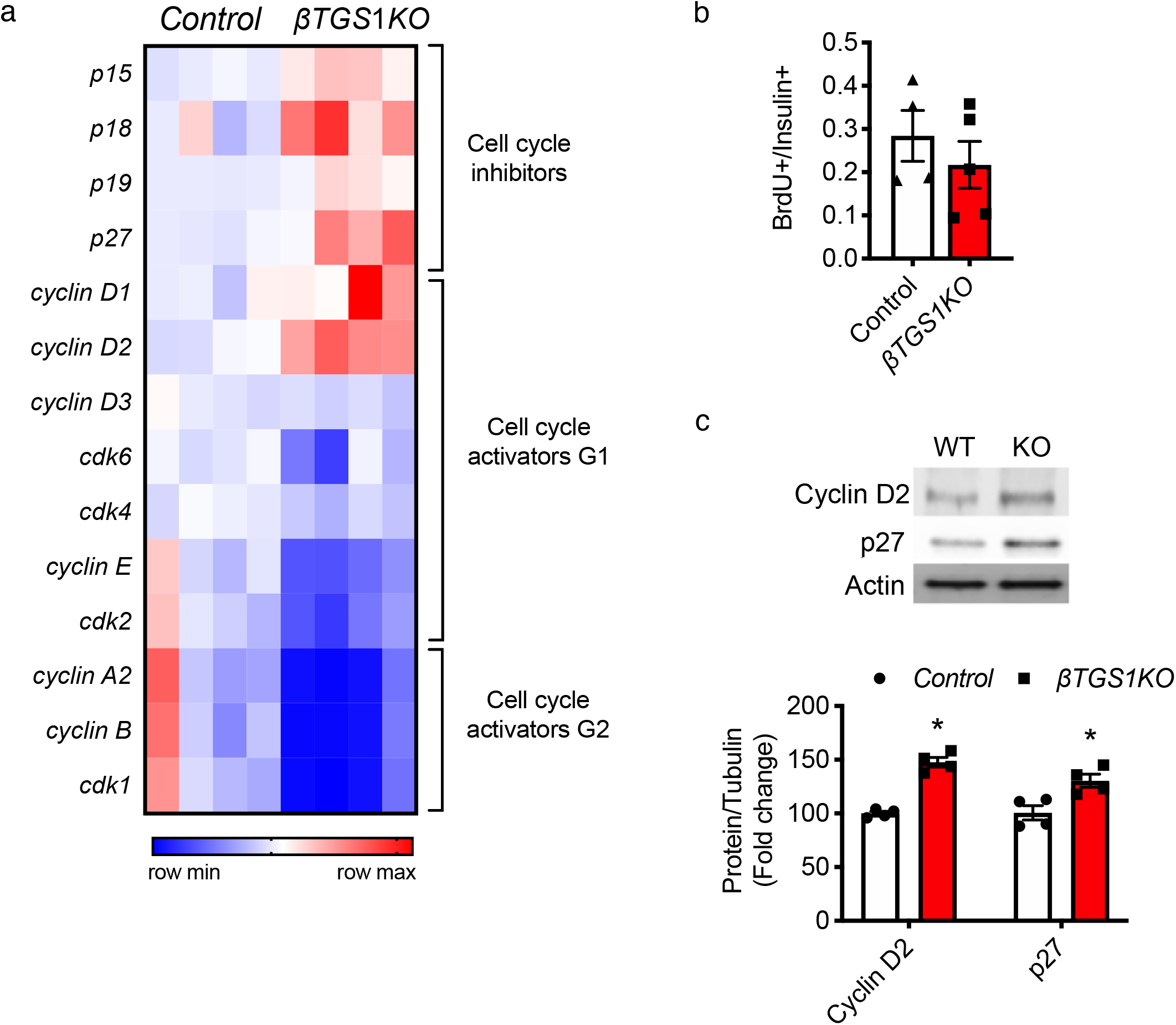
*β*-*cells with deletion of TGS1* exhibit cell cycle arrest. (a) Gene-expression heatmap of the differentially expressed genes related to cell cycle inhibitors, cell cycle components related to G1 phase and cell cycle components related to G2 phase from the RNA-Seq of *βTGS1KO* islets compared to control. Genes are represented in rows and mice in columns. (b) Assessments of proliferation by BrdU staining by FACS. (c) Immunoblotting and quantification for Cyclin D2, p27 and actin (n=4). Data expressed as means±s.e.m., \**p*<0.05. Nonparametric *U*-test (Mann–Whitney).

### TGS1 is important for β-cell adaptation to diet induced insulin resistance

Immunostaining of TGS1 of pancreas sections demonstrated an increase in TGS1 staining in β-cells from mice exposed to HFD for 8 weeks and T2D donors in comparison to controls (Figs. 1d and f). However, it is unclear if the increase in TGS1 in these models of diabetes plays a beneficial compensatory role or if the high levels observed have a negative impact and contribute to β-cells demise. We demonstrate that deletion of TGS1 in β-cell decreases β-cell mass and impairs glucose homeostasis. Interestingly, mice with heterozygous deletion of TGS1 in β-cells (*βTGS1^Het^)* exhibit normal glucose homeostasis on regular chow (Figs. 7a-b). Further, to test that aging was not affecting glucose homeostasis in these mice, glucose tolerance tests were performed in 1-year-old mice which showed no differences (Fig. 7c). To evaluate the adaptation of the β-cell mass and glucose homeostasis in response to HFD, *βTGS1^Het^* and controls (*RIP-Cre* and *TGS1^f/+^*) were fed with HFD for 8 weeks. TGS1 nuclear staining was increased in control mice as shown above (Figs. 1d and f). In contrast, staining of pancreatic sections from *βTGS1^Het^* mice shows a decrease in nuclear TGS1 staining (Fig. 7d, right). Glucose measurements during HFD administration show that *βTGS1^Het^* mice exhibit normal glucose during the first 2 weeks (Fig. 7e). However, *βTGS1^Het^* exhibit higher glucose levels at 8 weeks of HFD (Fig. 7e). *βTGS1^Het^* mice exhibited impaired glucose tolerance after 4 weeks of HFD (Figs. 7a, e-f). Surprisingly, β-cell mass was similar in *βTGS1^Het^* and control mice (Fig. 7g). These studies are consistent with the idea that TGS1 levels are important for the adaptation to HFD by decreasing insulin secretion.

**Figure 7.**
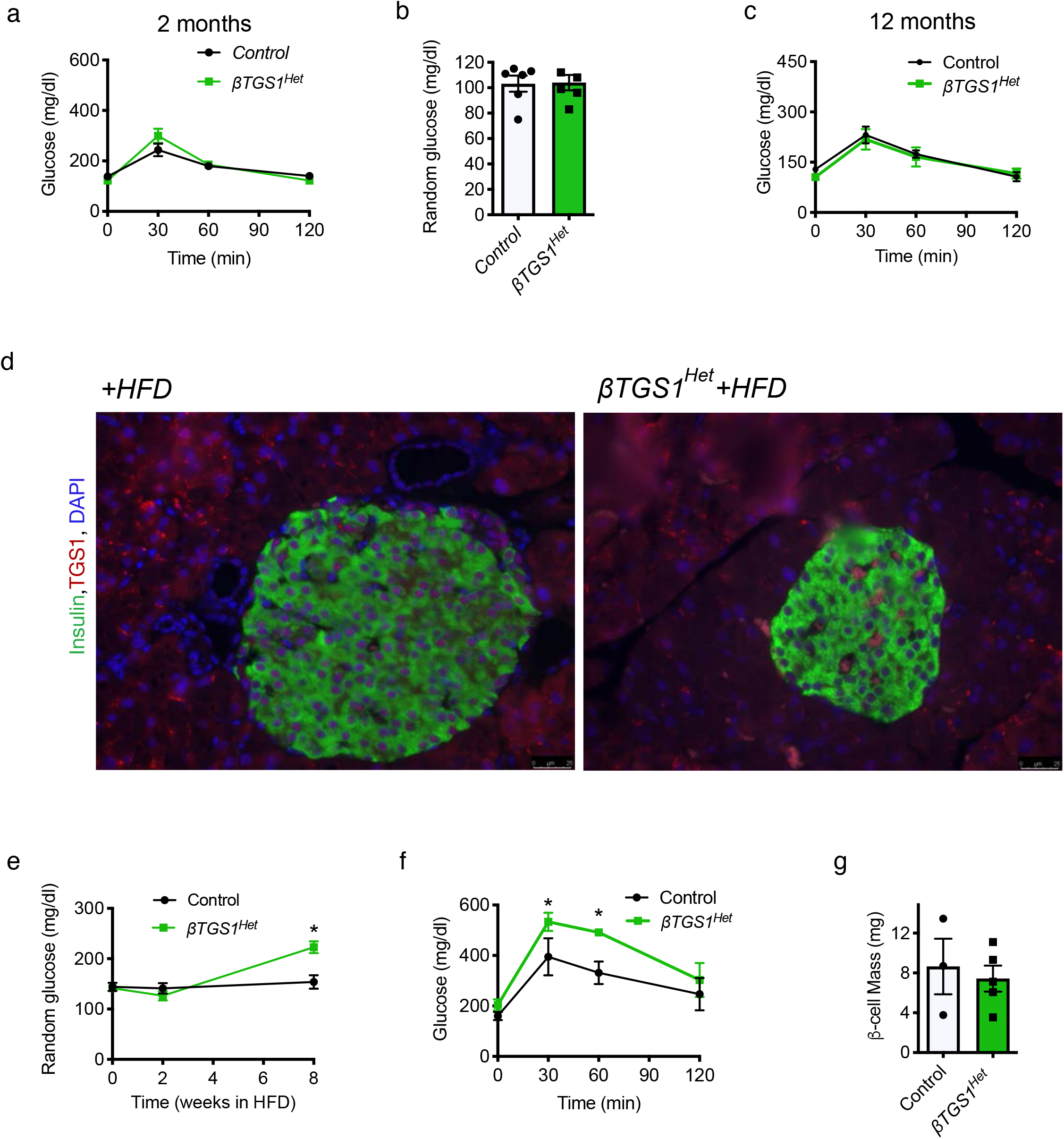
TGS1 heterozygous mice exposed to HFD exhibit impaired β-cell function without changes in β-cell mass. (a) IPGTT before HFD at 2 months of age. (b) Random fed glucose levels at 2 months of age. (c) IPGTT at 12 months of age. (d) Immunostaining of insulin (green), TGS1 (red) and DAPI (blue) in control and *βTGS1^Het^* and control mice fed HFD (n=4). (e) Random fed glucose levels during 8 weeks in HFD. (f) IPGTT after 4 weeks in HFD. (g) Assessment of β-cell mass. Data expressed as mean±s.e.m, \**P*<0.05 by nonparametric *U*-test (Mann–Whitney).

## DISCUSSION

The current studies assessed the role of TGS1 in β-cells. We showed that TGS1 is upregulated in β-cells in mouse models and human islets from T2D. The importance of TGS1 was further assessed in mice with constitutive and inducible inactivation of this gene in β-cells. These studies showed that TGS1 inactivation in mouse β-cells results in hyperglycemia and glucose intolerance as a result of a reduction in β-cells mass. We further identified TGS1 as a major regulator of β-cells survival and cell cycle progression. Finally, we also show that TGS1 plays an important function in insulin secretion. In conclusion, the current work identifies TGS1 as a major regulator of β-cells mass and function and demonstrates that this enzyme is important for adaptation to insulin resistance.

The current experiments showed that *βTGS1KO* mice maintain glucose homeostasis during the first month of life and slowly develop hyperglycemia (Fig. 3a). We found that reduced β-cell mass played a major role in the alteration in glucose homeostasis in these mice. The defect in β-cell mass was in part explained by induced β-cell apoptosis. A major finding of these experiments was the identification of TGS1 in the regulation of the UPR and ER stress. This conclusion was supported by RNAseq studies and validated by immunoblotting and electron microscopy. Our data show an increase in mRNA of BIP (hspa5) a major regulator of UPR pathways. This was accompanied by mRNA expression for components of the PERK, ATF6, and IRE1α pathways. Electron microscopy showed dilated ER validating the changes in mRNA and protein. This effect of TGS1 in survival is consistent with the increase in TUNEL observed in TGS1 deficient blastocysts suggesting that TGS1 might alter transcription of critical survival genes (17). In line with our findings, a main target of ATF-4, ddit3/CHOP mRNA was increased in *βTGS1KO* (Fig. 5c).

The morphometric analysis suggests that the increase in proliferation is not sufficient to compensate indicating a disbalance between apoptosis and proliferation or alterations in cell cycle. The increase in Ki67 without changes in BrdU suggested to us that β-cell from *βTGS1KO* failed to progress to the S phase (Figs. 6b). These abnormalities in cell cycle profile were further dissected by RNA-Seq expression of cell cycle profiles. These studies show an increase in drivers of G1-S transition including Cyclin D1 and 2 and cell cycle inhibitors that regulate G1-S transition including p18, p19, and p27. These changes were accompanied by a decrease in cyclin D complex partners cdk6, cdk4, other G1-S transition genes, cyclin E and cdk2 and G2 and M phase components cyclin A, cyclin B and cdk1 mRNAs (Fig. 6a). These experiments suggest that β-cell with TGS1 deletion entered the cell cycle but did not progress through different phases of the cell cycle. An interesting possibility is that genes involved in cell cycle arrest can contribute to the abnormalities of cell cycle in *βTGS1KO* mice. CHOP is a very well-known marker of ER stress but also is a DNA-damage inducible gene as gadd34. These genes were increased in the RNA-Seq data from *βTGS1KO* islets (Fig. 5c) and could and synergistically suppress cell growth as described (30). Further, another mRNA of genes related to growth arrest and DNA-Damage-inducible (GADD) like sesn1 and sesn2 (Sestrin 1 and 2) (31–33), are also upregulated in *βTGS1KO* islets (Fig. 5c). This suggests that TGS1 can modulate cell cycle not only by regulating cell cycle machinery but also genes involved in cell cycle arrest. It is difficult to determine if the primary mechanism is decreased proliferation or increased apoptosis but it is possible that the compensatory increase in proliferation by reduced apoptosis and hyperglycemia is defective.

Previous work has uncovered some of the molecules and pathways controlling β-cells in T2D. The current work adds TGS1 as a novel regulator of β-cells from mouse models of hyperglycemia and in human islets from T2D. The mechanisms for the increase in TGS1 levels in models of diabetes are not completely identified but we show that insulin in a paracrine fashion could be involved. Our data suggest that the increase in TGS1 plays a beneficial adaptive function in the adaptation of β-cells in T2D. Further validation of this positive role of TGS1 is supported by the abnormal glucose homeostasis in *βTGS1*^Het^ mice exposed to HFD. Importantly, these studies suggest that TGS1 is important to maintain insulin secretion in states of insulin resistance (Fig. 7). The important role of TGS1 in insulin secretion was also validated in *MIP-Cre^ERT^-TGS1KO* after TMX injection (Fig. 3j-l) and *βTGS1KO* at 1 month of age with no changes in insulin content or β-cell mass (Fig. 3i-m). Future experiments will be needed to understand how TGS1 regulates insulin secretion.

Major open questions after these studies are how TGS1 regulates the UPR, cell cycle components and insulin secretion. Answer to these questions will require a significant effort but these studies provide a steppingstone to uncover novel mechanisms in β-cells. TGS1 regulates methylation of snRNA and snoRNA and this mediates the biogenesis of small nuclear and nucleolar ribonucleoproteins (RNPs) complexes by binding to the survival of motor neurons (SMN) complex (snRNPs) and NOP proteins (snoRNPs) in the cytoplasm and nucleus respectively. snRNPs and snoRNPs are major constituents of the spliceosome and contribute to restoring homeostasis to RNA metabolism upon recovery from the stress. Future experiments can be designed to examine the role of mRNA splicing, transcription, and ribosome production in the observed β-cell phenotypes. Altered snRNPs biogenesis has been associated with defects in CBs biogenesis leading to splicing defects (28). snoRNAs also alter mitochondrial metabolism, modulate GSIS and responses to oxidative stress in islets, thereby providing a link between TGS1 and insulin secretion (34). Future studies could be designed to assess how these mechanisms regulate insulin secretion, ER stress and β-cell proliferation.

Here, we identify TGS1 as a novel regulator of β-cell mass and function. Our results suggest that an increase in TGS1 levels could play a beneficial role in the adaptation of β-cell to insulin resistance and diabetes. Additionally, these studies show that TGS1 regulates ER stress and cell cycle progression controlling β-cell mass. Another important conclusion from these studies is that decreased levels of TGS1 in states of insulin resistance could decreased insulin secretion affecting glucose homeostasis. Together, these data support the concept that elevation of TGS1 levels in β-cell and insulin-sensitive tissues could be a common marker of hyperglycemia/insulin resistance in diabetes. These studies strongly demonstrate the importance of TGS1 levels in β-cells and suggest that controlling TGS1 levels could be a novel therapeutic target to control glucose levels in T2D.

## Research Design and Methods

### Animals and Treatments

*RIP-Cre,* and mice with targeted deletion of TGS1 have been previously described (13–15,17,18). Studies were performed on mice on C57BL6J background. Results of the experiments are shown for male mice, but phenotypes were validated in female mice. Ages are shown in figure legends. All animals were maintained on a 12h light–dark cycle. All the procedures were approved by University of Miami IACUC committee and performed in accordance with University of Miami Animal Care Policies and the GUIDE for the care and use of laboratory animals.

### Metabolic Studies

Blood glucose levels were determined from blood obtained from the tail vein using Contour glucometer (Bayer). Glucose tolerance tests were performed on animals fasted overnight by intraperitoneally injecting glucose (2mg/kg). Plasma insulin concentrations were determined using a Mouse ultrasensitive Insulin ELISA kit (ALPCO).

### Islets studies

After islet isolation (35), islets were maintained at 37⊡°C in an atmosphere containing 20% oxygen and 5% CO_2_. Insulin secretion from isolated islets was assessed by static incubation (35). Briefly, after overnight culture in RPMI containing 5mM glucose and 10% FBS, islets were pre-cultured for 1h in Krebs–Ringer medium containing 2mM glucose. Groups of 20 islets in quadruplicates were then incubated in Krebs–Ringer medium containing 2 or 16mM glucose for 40min. Secreted insulin in the supernatant and insulin content was then measured using Mouse Ultrasensitive Insulin ELISA kit (ALPCO Immunoassays) and normalized to insulin total content.

### Immunoblotting

Islets from an individual mouse (150–300 islets) were lysed in lysis buffer (125□mM Tris, pH 7, 2% SDS and 1□mM dithiothreitol) containing a protease inhibitor cocktail (Roche Diagnostics). Protein quantity was measured by a bicinchoninic acid assay method, and 20-40μg of protein were loaded on SDS–PAGE gels and separated by electrophoresis. Separated proteins were transferred onto PVDF membranes (Millipore, Bedford, MA) overnight. After blocking for 1h in Intercept Blocking Buffer from LI-COR (Lincoln, NE), membranes were incubated overnight at 4°C with a primary antibody diluted in the same buffer followed by 1h incubation at room temperature with secondary antibodies from the same company. Antibodies used for immunoblotting are included in Supplementary Table 1, and membranes were developed using LI-COR Odyssey FC. Band densitometry was determined by measuring pixel intensity using NIH Image J software/Fiji (v2.1.0/1.53c (ref. (36)) freely available at http://rsb.info.nih.gov/ij/index.html) and normalized to tubulin, actin or total protein in the same membrane. Images have been cropped for presentation. Full-size images for the most important western blots are available from the authors on request.

### FACS

After overnight culture in RPMI containing 5□mM glucose, islets were dispersed into a single-cell suspension and fixed with BD Pharmingen Transcription Factor Phospho Buffer Set (BD Biosciences). Dispersed cells were incubated overnight with conjugated antibodies at 4°C. Dead cells were excluded by Ghost Dye Red 780 (Tonbo), and signal intensity from single stained cells and GFP was analyzed by mean fluorescent intensity in insulin-positive cells using BD LSR II (BD Biosciences). Antibodies used are included in Supplementary Table 1.

### RNA-Seq library preparation, sequencing, and data analysis

This data has been deposited in the GEO repository (pending). RNA Quality Control and DNase

#### Treatment

A total of 8 mice islets RNA samples were submitted to Ocean Ridge Biosciences (Deerfield Beach, FL) for mRNA-Sequencing. Total RNA was quantified by O.D. measurement and assessed for quality on a 1% agarose – 2% formaldehyde RNA Quality Control (QC) gel. The RNA was then digested with RNase free DNase I (Epicentre; Part # D9905K) and re-purified using Agencourt RNAClean XP beads (Beckman Coulter; Part # A63987). The newly digested RNA samples were then quantified by O.D. measurement. The newly digested RNA samples were then quantified by O.D. measurement and checked for quality.

#### Library Preparation

Amplified cDNA libraries suitable for sequencing were prepared from 250 nanograms (ng) of DNA-free total RNA using the TruSeq Stranded mRNA Library Prep (Illumina Inc.; Part # 20020595). The quality and size distribution of the amplified libraries were determined by chip-based capillary electrophoresis (Bioanalyzer 2100, Agilent Technologies). Libraries were quantified using the KAPA Library Quantification Kit (Kapa Biosystems, Boston, MA).

#### Sequencing

The 8 libraries were pooled at equimolar concentrations and sequenced in a total of 3 runs on the Illumina NextSeq 500 sequencer using two Mid Output v2 150 cycle kits (part# FC-404-2001) and one High Output v2.5 150 cycle kit (part# 20024907). In each case the libraries were sequenced with 76 nt paired-end reads plus 8 nt dual-index reads on the instrument running NextSeq Control Software version 2.2.0.4. Real time image analysis and base calling were performed on the instrument using the Real-Time Analysis (RTA) software version 2.4.11. Generation of FASTQ files: Base calls from the NextSeq 500 RTA were converted to sequencing reads in FASTQ format using Illumina’s bcl2fastq program v2.17.1.14 with default settings. Sequencing adapters were not trimmed in this step.

Gene ontology analysis was performed using EnrichR (29). GO Function process output comma-delimited (CSV) files were saved, and all those pathways having an adjusted p-value <0.05 were selected in the analysis. We selected commonly and highly ranked GO pathways, highly enriched in genes derived from the differential gene expression analysis.

### Immunofluorescence and morphometry

Formalin-fixed pancreatic tissues were embedded in paraffin and sectioned. Immunofluorescence staining was performed using primary antibodies described on Supplementary Table 1. Fluorescent images were acquired using a microscope (Leica DM5500B) with a motorized stage using a camera (Leica Microsystems, DFC360FX), interfaced with the OASIS-blue PCI controller and controlled by the Leica Application Suite X (LAS X). β-cell ratio assessment was calculated by measuring insulin and acinar areas using Adobe Photoshop in five insulin-stained sections (5μm) that were 200μm apart. To calculate β-cell mass, β-cell to acinar ratio was then multiplied by the pancreas weight. Assessment of proliferation was performed in insulin- and Ki67-stained sections and included at least 3,000 cells per animal. Apoptosis was determined using TUNEL assay (ApopTag Red in Situ Apoptosis Detection Kit, Chemicon) in insulin-stained sections. At least 3,000 β-cells were counted for each animal. Cell size was determined by immunostaining sections measuring the areas of individual β-cells from different experimental groups using NIH Image J software/Fiji. For dispersed cell staining, islets were gently dispersed after 5□min incubation with trypsin– EDTA (0.25% trypsin and 1□mM EDTA) in Hanks’ balanced salt solution without Ca^2+^ and Mg^2+^ (Gibco Invitrogen) at 37°C followed by fixation in 4% methanol-free formaldehyde onto poly-l-lysine-coated slides. All the morphologic measurements were performed in blinded manner.

### Electron microscopy

Ultrastructural characterization by transmission electron microscopy was performed after overnight culture (in RPMI containing 5mM glucose at 37°C). Islets were then fixed with 2% glutaraldehyde and then dehydrated and embedded in Epon by the Transmission Electron Microscopy Core at University of Miami. Ultrathin sections were stained with uranyl acetate and lead citrate, and images were recorded digitally using a Philips CM-100 electron microscope.

### Human organ donors

Human pancreas tissue samples (from the head of the pancreas, non-diabetic individuals and T2 diabetic individuals (n=4), male and female, ages=15-52 years-old) from the Human Islet Cell Processing Facility at the Diabetes Research Institute, University of Miami.

### Statistical analysis

Data are presented as mean□±□SEM. Student t test was employed to assess statistical difference between means of two groups in one time point. The statistical significance of differences between the various conditions was determined by nonparametric *U*-test (Mann– Whitney) using Prism version 9 (GraphPad Software, San Diego, CA). Results were considered statistically significant when the p value was equal or less than 0.05.

## Data availability

All relevant data are available from the authors on request.

## CRediT author statement

M.B.-R., P.R.Ll. and A. L. performed the experiments and analyzed results. M.B.-R. and E.B.-M. wrote the article and designed the experiments. J.K.R. generated mice. M.B.-R. and E.B.-M. contributed to discussion and reviewed and edited the manuscript. M.B.-R. is the guarantor of this work and, as such, had full access to all the data in the study and takes responsibility for the integrity of the data and the accuracy of the data analysis.

---

**Supplemental Table 1.** Antibodies

| Antibody | Specie | Source | Catalog | Concentration |
| --- | --- | --- | --- | --- |
| Actin | Mouse | Sigma | A5441 | 1:4000 |
| Active caspase 3 BV650 | Rabbit | BD Trans. | 564096 | 5 µl per test |
| ATF4 | Rabbit | Santa Cruz | 200 | 1:500 |
| BrdU | Mouse | Cell Signaling | 5292 | 1:200 |
| Coilin | Rabbit | Cell Signaling | 14168T | 1:400 |
| Cyclin D2 | Mouse | Santa Cruz | 593 | 1:100 |
| Insulin | Guinea Pig | Dako | A0564 | 1:400 |
| Insulin APC-Conjugated | Rat | R&D Systems | IC1417A | 1 µl per test |
| Ki67 | Rabbit | Vector | NCL-Ki67p/VPK451 | 1:200 |
| p-EIF2alpha | Rabbit | Cell Signaling | 3398 | 1:1000 |
| p27 | Mouse | BD Transduction |  | 1:203 |
| t-EIF2alpha | Rabbit | Cell Signaling | 5324 | 1:1000 |
| TGS1 | Rabbit | Bethyl Laboratories | 00467 | 1:250 (IHC and IF) |
| TGS1 | Rabbit | Bethyl Laboratories | A304-922A | 1:500 (WB) |
| Tubulin | Mouse | Sigma | T5168 | 1:3000 |
| U1 | Mouse | Santa Cruz | sc-390899 | 1:100 |
| XBP1 | Rabbit | Santa Cruz | 7160 | 1:100 |

